# Ocular Dominance Plasticity: Measurement Reliability and Variability

**DOI:** 10.1101/2020.07.27.211144

**Authors:** Seung Hyun Min, Ling Gong, Alex S. Baldwin, Alexandre Reynaud, Zhifen He, Jiawei Zhou, Robert F. Hess

**Author notes:** Correspondence can be achieved via.

## Abstract

In the last decade, studies have shown that short-term monocular deprivation strengthens the deprived eye’s contribution to binocular vision. However, the magnitude of the change in eye dominance after monocular deprivation (i.e., the patching effect) has been found to be different between for different methods and within the same method. There are three possible explanations for the discrepancy. First, the mechanisms underlying the patching effect that are probed by different measurement tasks might exist at different neural sites. Second, test-retest variability in the measurement might have led to inconsistencies, even within the same method. Third, the patching effect itself in the same subject might fluctuate across separate days or experimental sessions. To explore these possibilities, we assessed the test-retest reliability of the three most commonly used tasks (binocular rivalry, binocular combination, and dichoptic masking) and the repeatability of the shift in eye dominance after short-term monocular deprivation for each of the task. Two variations for binocular phase combination were used, at one and many contrasts of the stimuli. Also, two variations of the dichoptic masking task was tested, in which the orientation of the mask grating was either horizontal or vertical. This makes five different measurement methods in all. We hope to resolve some of the inconsistencies reported in the literature concerning this form of visual plasticity. In this study, we also aim to recommend a measurement method that will allow us to better understand its physiological basis and the underpinning of visual disorders.

## 1 Introduction

Over the present decade, there has been increasing evidence that a new form of temporary binocular plasticity exists in human adults. For instance, patching an eye for a short period strengthens that eye’s contribution to binocular vision [1, 2]. This has been demonstrated for patching periods as short as 15 minutes [3, 4]. Here, we will refer to this neuroplastic change in ocular dominance as a result of short-term monocular deprivation as *the patching effect*. The patching effect lasts for 30-90 minutes, and maybe beyond even that time [5, 1, 4]. It can be induced by both opaque and translucent patches, and by dichoptic video presentation that emulates the effects of patching one eye [6, 7]. The patching effect has been demonstrated with psychophysical, electrophysiological [8, 9] and neuroimaging [10, 11, 12] measurements. The change in sensory eye dominance as a result of short-term patching seems to be reciprocal between the eyes: the contrast gain of the patched eye is enhanced and that of the non-patched eye weakened [13, 2].

Patching studies agree that the patched eye strengthens after patching. However, the magnitude of the patching effect has been found to vary. Inconsistent results have been found between different measurement methods. There are three possible explanations for the discrepancy. First, the patching effect might be a complex phenomenon, rather than a change in a single factor (like an increase in one eye’s input gain). In other words, the mechanisms underlying the patching effect that are probed by different measurement tasks might exist at different neural sites. For example, removal of phase information induces the patching effect if it is measured with a binocular rivalry task [6] but not if it is measured with a binocular combination task [6, 7]. Moreover, the patching effect has been shown to be of larger and longer lasting in the chromatic visual pathway than in the achromatic visual pathway if it is measured with binocular rivalry [14] but not if it is measured with binocular combination [15]. Furthermore, the site of action has been argued to be at an early stage (i.e. striate) in cortical processing by some [16, 17, 7] and at a later stage (i.e. extra-striate) by others [6, 3, 18]. Second, test-retest variability in the measurement might have led to inconsistencies, even within the same method [5, 19]. It might be larger for some tests than for others. Third, the patching effect itself in the same subject might fluctuate across separate days or experimental sessions. This possibility has not been explored but may be important for some testing protocols.

Most previous studies have measured the effect of short-term patching once for each subject and experimental condition. This practice assumes that the respective psychophysical methodology is reliable and that the patching effect is stable across days for each subject. In this study, we question these assumptions. We undertook an experiment using each task across two experimental sessions. The test-retest reliability of the three most commonly used tasks (binocular rivalry, binocular combination, and dichoptic masking) and the repeatability of the patching effect for each of the task were evaluated. Two variations for binocular phase combination were used, at one [2] and many contrasts of the stimuli [4]. Also, two variations of the dichoptic masking task were tested, in which the orientation of the mask grating was either horizontal or vertical [20]. This makes five different measurement methods in all. We hope to resolve some of the inconsistencies reported in the literature concerning this form of visual plasticity. We will also aim to recommend a measurement method that will allow us to better understand its physiological basis and the underpinning of visual disorders. To do so, we assessed four properties of each task:

1. Baseline reliability: how well is the baseline performance (i.e., no patching) correlated for each subject between days?
2. Patching effect reliability: How well is the magnitude of the patching effect correlated for each subject between days?
3. Baseline measurement variability: What is the expected measurement variability from the task alone, and how does this compare to the overall variability in the baseline conditions?
4. Patching effect measurement variability: What is the expected measurement variability from the task alone in the patched conditions, and how does this compare to the overall?

## 2 Materials and Methods

### 2.1 Subjects

We used data from 88 adults (age range = 18-33) with normal or corrected-to-normal vision in this study. The data of 62 subjects have already been reported in publications [20, 4, 21, 5]. For this study alone, we tested 26 additional subjects. Some subjects completed more than one condition. *Therefore, the total number of data points in this study amounts to 148, each of which represents a subject who performed a unique experiment for two experimental sessions*. This study adhered to the Declaration of Helsinki and was approved by the Institutional Review Boards at McGill University and Wenzhou Medical University. All subjects provided informed written consent. All subjects performed each experiment for two sessions that were separated by at least 24 hours.

### 2.2 Monocular Deprivation

In all experiments, the dominant eye of the subject was patched. The eye dominance was determined by the Miles test [22]. For some psychophysical tasks we tested different patching durations (ranging from 15 to 180 minutes). Subjects performed each experimental session twice (i.e., same patching duration) on separate days. A translucent patch was used. It deprives all form information and reduces the luminance for the patch eye by 20%. During patching, subjects either browsed the web with their computer or phone. We were only interested in the immediate patching effect. We did not analyse the decay in the effect over the subsequent hours. Therefore, only data that were obtained immediately after patch removal (within 10 minutes) were included in our analysis.

### 2.3 Psychophysical Tasks

In this study, we investigate four psychophysical tasks (i.e., five variations in total). Each task is described in detail in this section. Moreover, we extracted a subset of data from four published studies [20, 5, 4, 21]. In this section, we elaborate on the rationale for the data extraction, the process of data analysis, and the experimental procedure for each psychophysical method.

#### 2.3.1 Binocular Rivalry

In this method, conflicting stimuli are shown to the two eyes. The relative strength of each eye is assessed by measuring the length of time for which each eye suppresses the other. Data from 30 subjects were collected for a previous study [5]. We reused the baseline measurements from Finn et al (2019). An additional 15 subjects were then tested as part of the current study. Therefore, data from 45 subjects were included in the binocular rivalry analysis. The apparatus for the experiment in the previously published study and the details of the methodology are explained in Finn et al (2019) [5]. The apparatus for the testing performed on the additional 15 subjects is described in this paper.

##### 2.3.1.1 Stimuli

In the binocular rivalry task, two oblique Gabor gratings at +45° and −45° were shown separately to the two eyes. They gratings had a spatial frequency of 1.5 c/deg, a spatial sigma of 1.3 degrees of visual angle and a contrast of 50%. Shutter glasses were used for the stimulus presentation. Each testing block lasted 3 minutes. Subjects reported continuously using the keyboard whether they perceived a left oblique grating, right oblique grating, or mixed percept throughout the test.

##### 2.3.1.2 Procedure

In the study of Finn et al. (2019), the patching effect was compared between two experimental conditions [5]. This was to test whether exercise during the patching period enhanced the patching effect. Because the patching conditions were not identical, we cannot use them for the analysis we are conducting in this study. The baseline measurements made on the two testing days were identical, however. Therefore, we excluded data from post-patching measurement, but included the baseline data to measure the test-retest variability of binocular rivalry.

However, since we were interested in evaluating the repeatability of the patching effect as measured in binocular rivalry, we collected more data. To do so, we tested 15 additional subjects for this study. The subjects first performed the baseline measurement in which the binocular rivalry task was performed four times (Figure 1). The binocular rivalry task was interleaved with a binocular combination task (the data from the combination task were not used for analysis). This was to make the procedure here more comparable with that used to compare two forms of the combination task (as described in the next section). The baseline tests therefore consisted of four experimental blocks of binocular combination and binocular rivalry tasks. After patching for 120 minutes, the subjects were tested again using binocular combination and binocular rivalry for two experimental blocks (two blocks per task).

**Figure 1:**
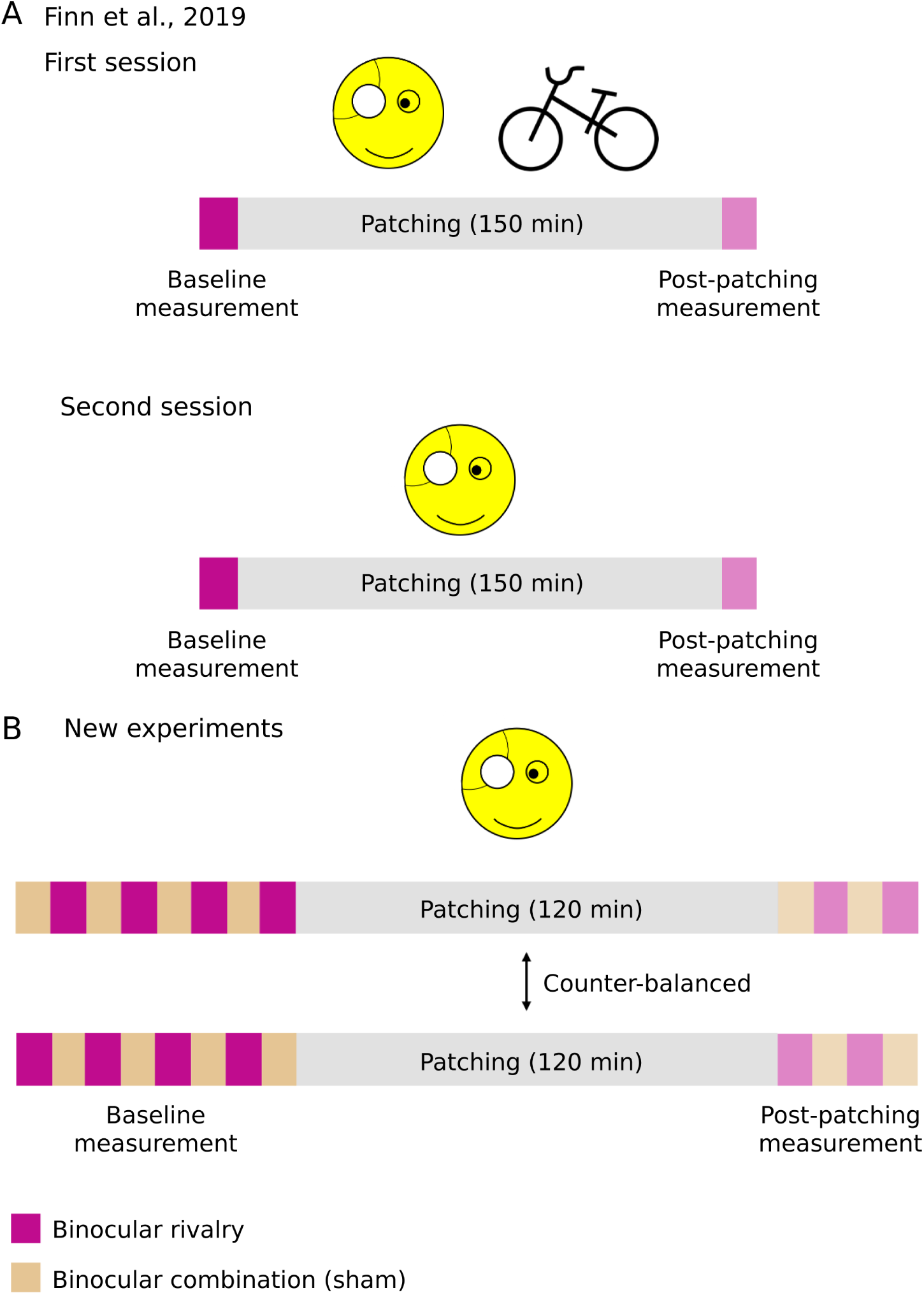
Procedures of experiments using binocular rivalry. A) Procedure of the experiment in the study of Finn et al. (2019). B) Procedure of the new experiments in our study.

##### 2.3.1.3 Data Analysis

We assigned each Gabor’s orientation to the role of each eye’s contribution in perceptual dominance. We computed the ocular dominance index (ODI) as follows:

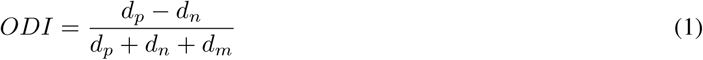

where d_*p*_, d_*n*_ and d_*m*_ are the total response durations of the percept perceived by the patched eye, non-patched eye and both eyes (i.e., mixed percept), respectively. When ODI is positive, the total response duration for the percept perceived by the patched eye is longer than that for the non-patched eye’s percept. When ODI is negative, the total response duration for the percept perceived by the non-patched eye is longer than that by the patched eye.

#### 2.3.2 Binocular Phase Combination at One Contrast

In this task, the subject adjusts dichoptically presented gratings so that the fused percept indicates that the contribution from the two eyes is balanced [23]. New data were collected from 15 subjects. Details on this task can be found in Zhou et al. (2013)[2] and Zhou et al. (2017)[24].

##### 2.3.2.1 Stimuli

Two separate horizontal sine-wave gratings (0.46 cycle/°, 4.33°*×*4.33^°^) with equal and opposite phase shifts (+22.5° and −22.5°) relative to the center of the screen were presented to the two eyes. The perceived phase of fused stimuli would be 0 if the two eyes contributed equally to binocular fusion (see Figure 1). The subjects were asked to locate their perceived middle portion of the dark patch in the fused grating by positioning a flanking 1-pixel reference line. The stimuli were displayed until subjects completed the tasks. The contrast of the stimuli shown to the non-dominant eye (i.e., non-patched eye) was set at 100% for each subject. Moreover, the contrast of the stimuli shown to the dominant eye (i.e., patched eye) was set so that both eyes contributed equally to binocular vision (i.e., binocularly perceived phase = 0). The contrast of the stimuli shown to the non-dominant eye was not uniform across subjects. Therefore, there was only one contrast ratio between the stimuli shown separately to the eyes for every subject.

##### 2.3.2.2 Procedure

The experimental protocol is identical to the interleaved design described in Section 3.1.2. The subjects performed baseline measurements with psychophysical tasks of binocular combination at one and multiple contrasts (another variation of binocular phase combination, described below in Section 2.3.3). They completed four experiment blocks of the two different binocular combination tasks (four blocks per task). Then they were patched for 120 minutes. During patching, they performed tasks such as reading and web browsing. After patching, they were tested again using the two methods of binocular combinations for two experimental blocks (Figure 2). We randomized the order of the task to be tested and maintained the order across two experimental sessions for each subject.

**Figure 2:**
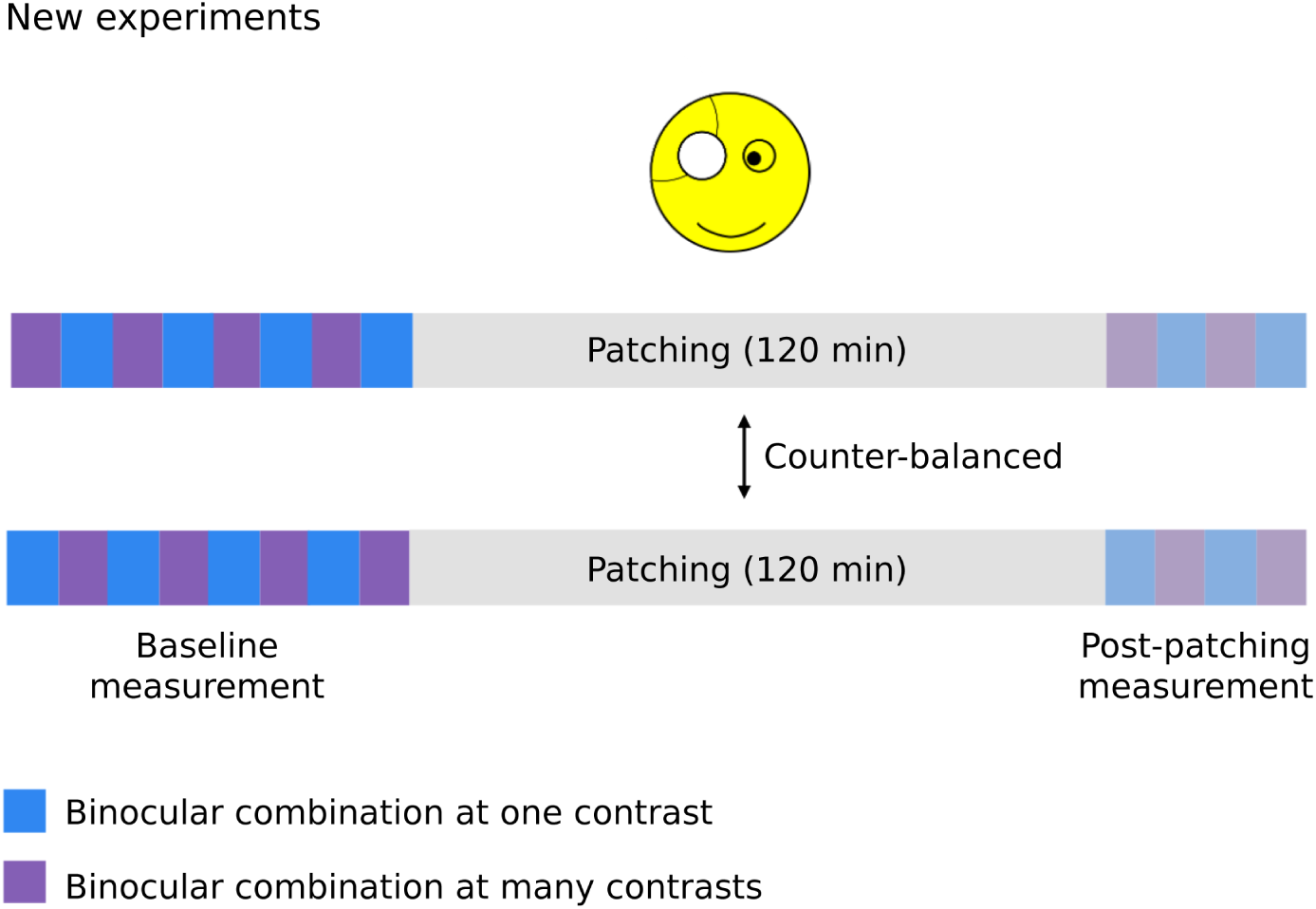
Procedure of the new experiments using, but not limited to, binocular combination at one contrast.

#### 2.3.3 Binocular Phase Combination at Many Contrasts

In this task, the percept of the subject is indicated when fusing dichoptic gratings at a range of different dichoptic contrast ratios. This allows one to calculate the interocular contrast ratio at which the percept would indicate a balanced input. Data from 19 subjects have already been collected in previous studies [4, 21]. Several subjects from this cohort were patched for more than one duration. Moreover, we tested 15 more subjects to directly compare the test-retest repeatability from this task with that from the binocular phase combination at one contrast. In sum, there were 60 unique data points, each of which is one subject being patched for a particular duration. The equipment and methodology are explained in detail in the previous studies [4, 21]. Only the apparatus that were used to collect the data of the 15 new subjects are described in Section 4 of Methods.

##### 2.3.3.1 Stimuli

The stimuli are very similar to those in binocular combination at one contrast. Two slightly offset horizontal sinusoidal gratings were presented to the two eyes. The phase difference was 45°: +22.5° for one eye and −22.5° for the other eye. If the two eyes contribute equally to binocular vision, the fused phase percept will appear as exactly the average of the two gratings phases. This is equivalent to the perceived phase of zero (see Figure 1).

The interocular contrast ratio between the eyes was changed by increasing the contrast of one eye’s stimulus while decreasing the contrast of the other eye’s stimulus (see Figure 1). Then, the interocular contrast ratio at a perceived phase of 0 degrees was estimated using a contrast gain model [23]. By comparing the binocular balance before and after patching, we calculated the shift in ocular dominance.

We set five interocular contrast ratios 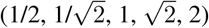 for baseline measurement, and three for post-patching measurement 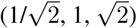. This was designed to shorten the test durations to as short as 3 minutes in the post-patching, measure. In the binocular phase combination at one contrast task (Section 2), only a single ratio (1) was used.

##### 2.3.3.2 Procedure

From the study of Min et al., 2018, we extracted data of 14 subjects who were patched for various durations (15 to 180 minutes). Prior to patching, the subjects performed the baseline experiments. After patching for an assigned duration, they completed post-patching experiments at several timepoints between 0 to 96 minutes after patching. All subjects performed each experimental session twice. Therefore, we were able to include data from baseline and post-patching assessments to evaluate the test-retest repeatability of the task. We only extracted post-patching data at the first three measured post-patching timepoints and averaged the values across time.

Data from 9 subjects were reused from the study of Min et al., 2019 [21]. One of the subjects from the study was excluded (as they also participated in the study of Min et al., 2018). The protocol is similar to the one described above, except that the subjects were patched for 120 minutes. Subjects performed the experiment for five consecutive days. We extracted data only from the first two days of the study. Post-patching data from 0 to 6 minutes were averaged to quantify the immediate patching effect.

As described in Section 2.2, we tested 15 more subjects to directly compare the test-retest repeatability between the two variations of binocular phase combination. The procedure is described in Section 2.2.

##### 2.3.3.3 Data Analysis

We averaged the perceived phases across two configurations from each subject. We then fitted these means of perceived phases into a contrast gain control model introduced by Ding and Sperling [23]:

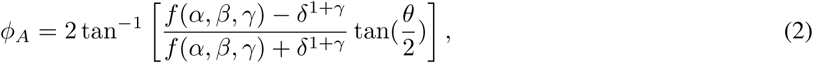

where

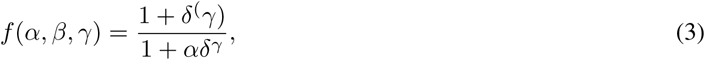

*ϕ*_*A*_ = perceived phase from the fused percept of two stimuli, *α* = gain factor which determines the contrast balance ratio when both eyes contribute equally to binocular vision, *γ* = slope of the function when both eyes contribute equally to binocular vision, *θ* = fixed phase displacement between eyes (45°), *δ* = interocular contrast balance ratio. After we fitted our data to the contrast gain model function [23], we estimated the two free parameters *α* and *γ*. We bootstrapped responses trial-to-trial and generated each measurement’s sample of values to generate standard errors for each data point.

*α*_*ratio*_ = contrast balance ratio when both eyes contribute equally to binocular vision in linear scale, *α*_*dB*_ = *α*_*ratio*_ in log scale. When the contrast shown to the dominant eye is as twice as strong as the non-dominant to reach the balance point (*α*_*DE*_ = 2*α*_*NDE*_), then *α*_*ratio*_, thereby resulting in *α*_*dB*_ = 6 dB.

We converted *α*_*ratio*_ into *α*_*dB*_ to avoid bias for the dominant eye when we quantify binocular balance. We normalized the contrast balance ratios by calculating for the differences in contrast balance ratios between baseline and after patching (dB). Therefore, when Δ contrast balance ratio = 0, it represents no change after patching. While a positive Δ contrast balance ratio indicates the shifting of ocular dominance favors the dominance eye (the patched eye).

#### 2.3.4 Dichoptic Masking Task

All data of 14 subjects for this experiment have already been used in a previous study [20]. No additional subjects were tested. The apparatus and details of the methodology are further explained in the previous study [20].

##### 2.3.4.1 Stimuli

One sinusoidal grating of 0.5 c/deg was presented to each eye. Gratings were presented in a circular raised-cosine envelope. The diameter was 5 degrees of visual angle. The temporal envelope for presenting the gratings was a Gabor (temporal frequency of 2 Hz, duration sigma 500 ms). The contrast in log units (dB) was computed as:

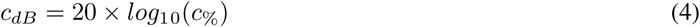

A contrast of 1% translates to 0 dB. A twofold threshold elevation from masking gives a 6 dB difference between detection thresholds with and without the mask.

The experiment used a two-interval forced choice procedure. Contrast detection thresholds were measured under three conditions: i) monocularly in the eye to be patched (no mask), ii) monocularly in the eye to be patched with a dichoptic mask grating shown to the other eye that had the same orientation as the target (parallel), iii) similar to ii), but with the mask having an orthogonal orientation (if the left eye’s grating were 45°, the right eye’s grating would be −45°). The mask contrast was fixed at 4%. When a mask was shown, it would be presented to the non-patched eye in both intervals. In only one of the intervals, the target grating would be shown (to the patched eye). The subject reported the interval (first or second) in which the target grating was presented.

##### 2.3.4.2 Procedure

Baldwin and Hess (2018) asked the subjects to perform baseline tests which measured the detection threshold of the patched eye, as well as the detection threshold of the patched eye when the mask grating was shown to the non-patched eye (i.e., masked threshold) in two different orientations (parallel and cross) [20]. Then the dominant eye was patched for 150 minutes during which the subjects performed tasks such as reading and web browsing. After patch removal, subjects were asked to immediately perform three blocks of post-patching measurements. The post-patch tests included three test blocks and measured masked threshold of the patched eye. The sequence of the testing block was either parallel-cross-parallel or cross-parallel-cross. All subjects completed both sequences in a randomized order across the two sessions. The sequence order of the post-test was counterbalanced because the shift in eye dominance after patching has been known to decay over time.

### 2.4 Apparatus for the New Experiments

We programmed the new experiments in MATLAB 2012a using PsychToolBox 3.0.9 [25, 26]. We presented the stimuli on a Mac computer with gamma-corrected head mounted goggles (NED Optics Groove pro, OLED). They had a refresh rate of 60 Hz and resolution of 1920 *×* 1080 to the screen for each eye. The maximum luminance of the goggles was 150 cd/m2.

### 2.5 Standardized Data Analysis

Data were analyzed using R and Python. We categorized our entire dataset into baseline data and those that quantify the magnitude of change in eye dominance after patching (i.e. patching effect). To investigate the four aspects set out in our introduction (baseline reliability, patching effect reliability, baseline measurement variability, and patched measurement variability), we standardized the raw score from each psychophysical task. We converted the raw data into z-score using this formula:

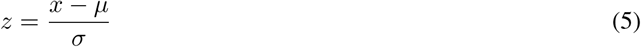

where *x* is the raw data, *μ* is the mean of the sample, *σ* is the standard deviation of the sample. A z-score indicates how far a data point is from the mean of a particular dataset. For example, if a data point has a z-score of 1.0, it is at one standard deviation from the mean.

The results from each task are analysed in a similar way. Below we describe each column of our figures.

#### 2.5.1 Column (i): Baseline and Patching Effect Reliabilities

To assess test-retest repeatability, Pearson’s correlation was calculated using raw data. A strong correlation indicates that a subject’s performance from the first session is a good predictor of that in the second session. In this column, figures also show the conversion of raw data into z-scores.

#### 2.5.2 Column (ii): Baseline and Patching Effect Measurement Variabilities

Throughout this paper, Bland-Altman plots [27] are used to evaluate the measurement variability of either baseline or the patching effect. We generated the Bland-Altman plots using the z-scores. Each test-retest pair is plotted as a single point. Its location on the y-axis is the difference between the z-scores from the first and second sessions. Its position on the x-axis is the mean z-score across the two sessions. The mean difference between the two days (across subjects) is indicated by the central horizontal dashed line. By computing 95% confidence interval limits of agreement (mean difference between sessions ± 1.96 SD), we calculated a measure of the test-retest variability (outer dashed lines).

Since the two experimental sessions were separated by at least 24 hours, we reasoned that the variability indicated by the outer dashed lines might arise from a combination of factors. The first of these is the measurement error which arises from the task design and testing procedure. The second would be day-to-day variability in the measured physiological mechanisms. We estimated the first of these factors by computing the expected standard error that arises solely from the psychophysical task of interest. To obtain a representative standard error for each task, the median of the standard error from each dataset of the task across two sessions was obtained. This was the standard error for a single measure, but as the Bland Altman plots analyse the difference between two measurements then the standard errors of both needed to be accounted for. We did so by multiplying the single standard error by 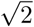. To convert this “difference standard error” to a 95% confidence interval it was multiplied by 1.96. We calculated the range between the mean of the differences between the two sessions and the expected 95% confidence interval from the measurements. We subsequently shaded this range in grey. This shaded grey region represents the expected measurement variability from the psychophysical task itself. Where the dashed lines indicating the limits of agreement are wider than this shaded region, then that represents an additional source of variability beyond the measurement alone.

#### 2.5.3 Column (iii): Baseline and Patching Effect Correlations

Finally, whether the performance of a single subject across days was significantly more correlated than a mismatched pair of subjects was evaluated. To do so, we plotted histograms of randomly-sampled values from the first session of one subject and second session of another randomly-selected subject 1000 times from the dataset of each task. Then, correlation coefficients were computed. These estimated correlation coefficients were then compared to the correlation coefficient between the two performance sessions of the same subject. If a correlation is robust, then it should deviate significantly from the histogram. These figures are shown in column (iii) in Results.

## 3 Results

### 3.1 Baseline Measurement

To assess the test-retest variability of the psychophysical tasks, we incorporated data from baseline measurement into our data analysis. Each subject performed two experimental sessions that were separated by at least 24 hours. Baseline results are shown in Figure 6.

**Figure 3:**
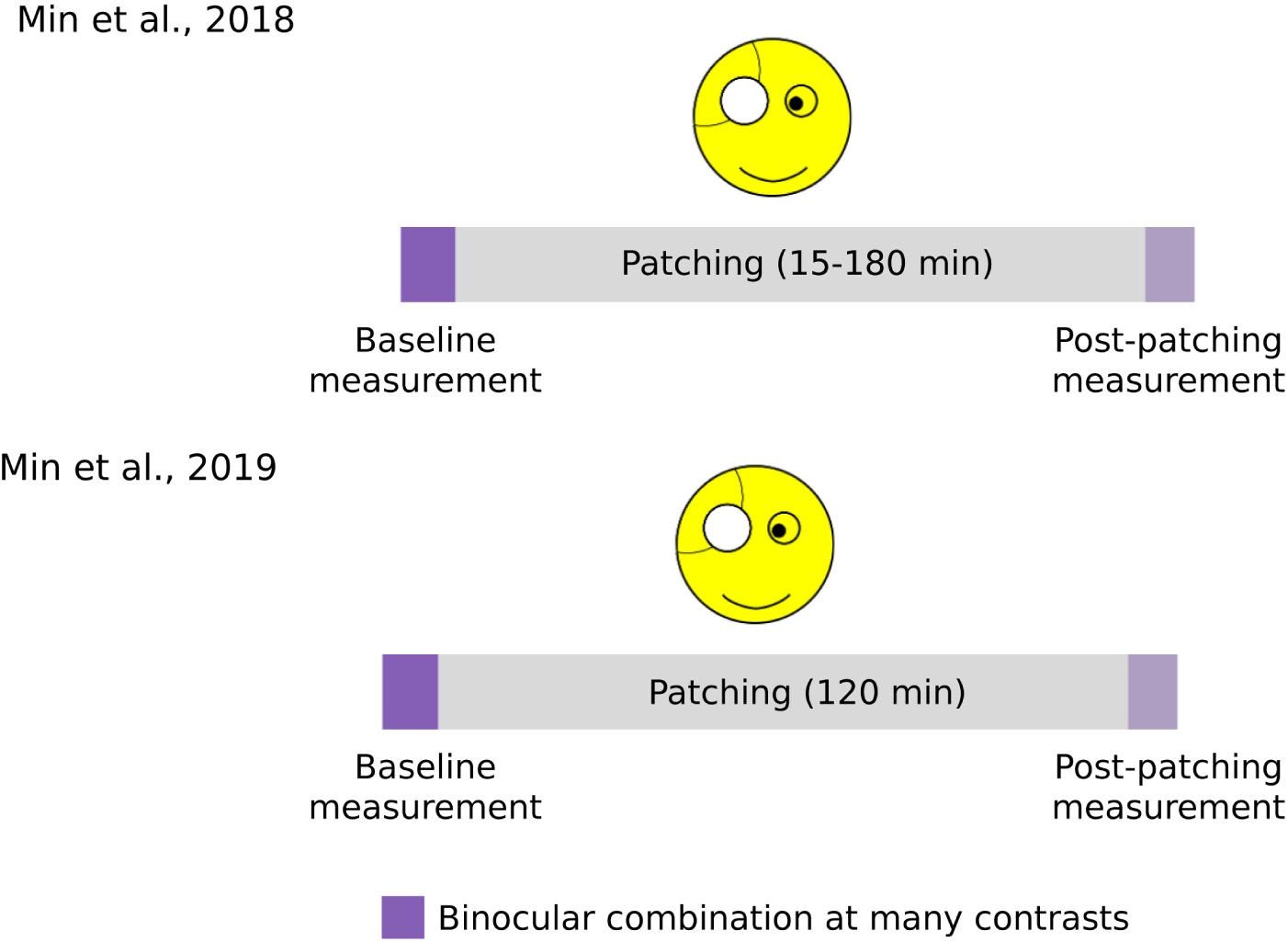
Procedure of experiments using binocular combination at many contrasts.

**Figure 4:**
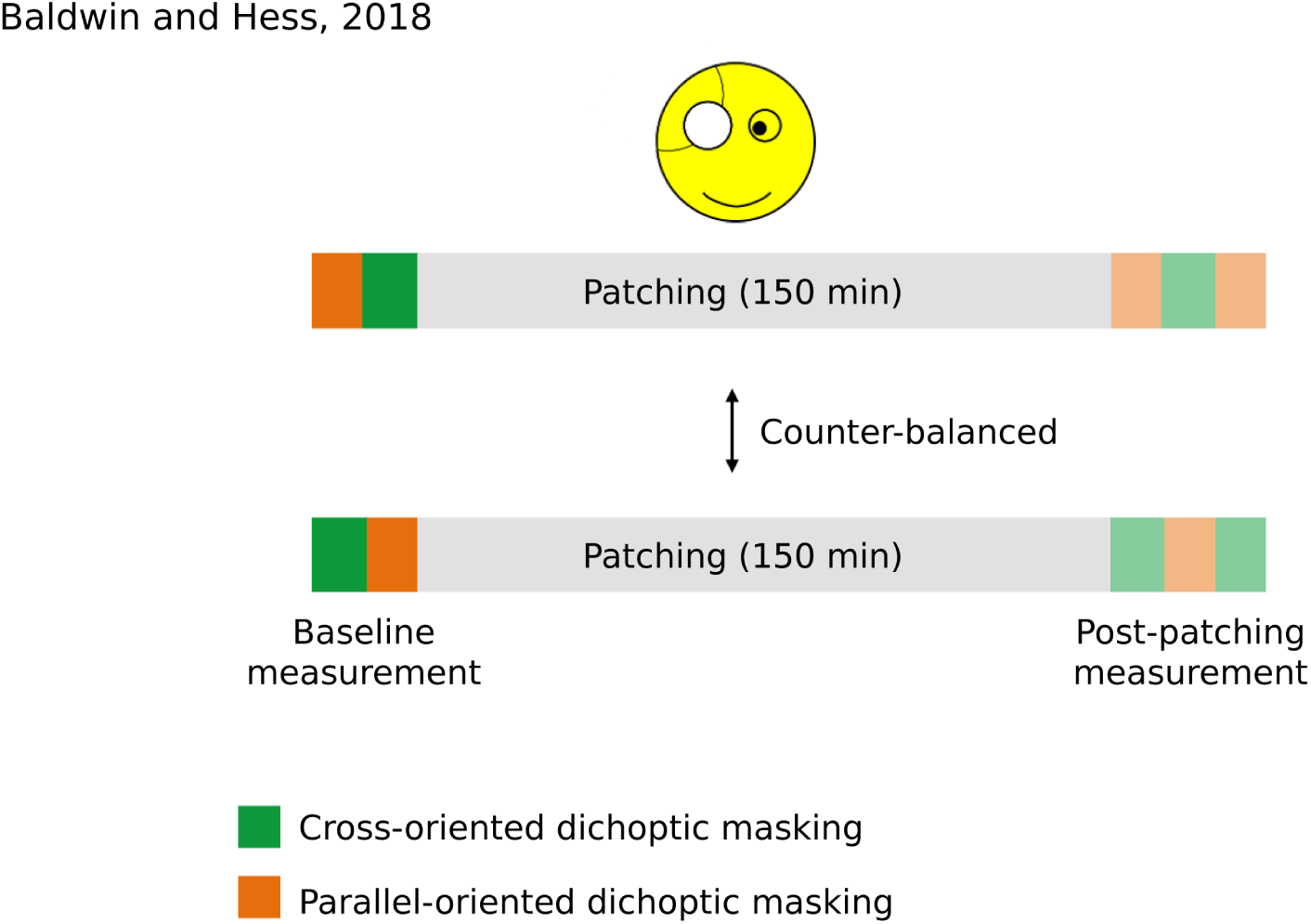
Procedure of experiments using dichoptic masking. The figure has been adapted from the previous study by Baldwin and Hess (2018) [20].

**Figure 5:**
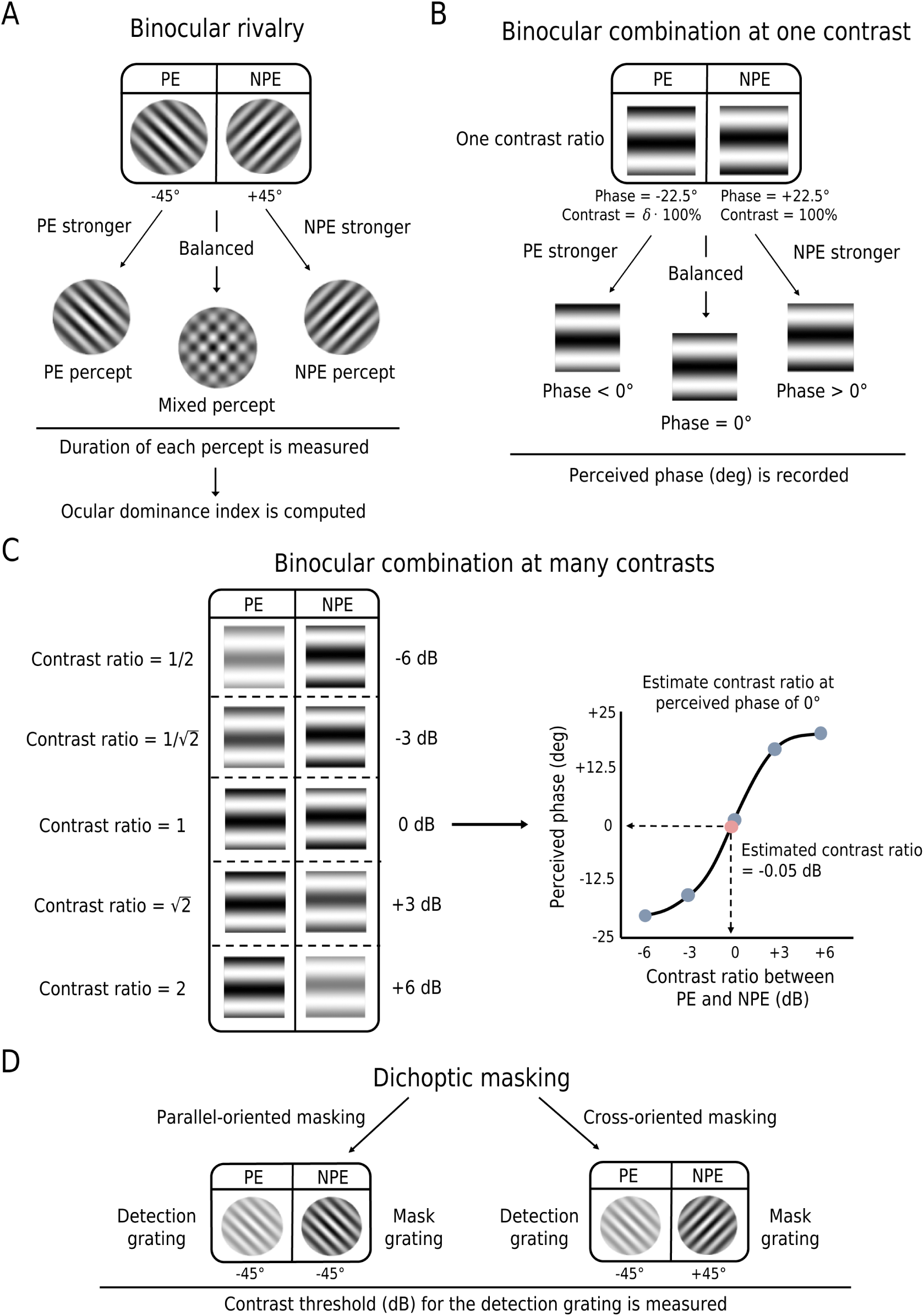
An illustration of stimuli in the five psychophysical task variations. PE = patched eye, NPE = non-patched eye. A) Binocular rivalry. Two gratings in different orientations are shown separately to both eyes. When the patched eye is dominant, the grating shown to the patched eye will dominate the conscious visual awareness. B) Binocular phase combination at one contrast. Two fusible gratings were shown dichoptically. Subjects were asked to locate using the keyboard the center of the darkest strip within the middle segment of the fused grating. C) Binocular combination at many contrasts. Two fusible gratings were shown separately to both eyes. Subjects were asked to locate using the keyboard the center of the darkest strip within the middle segment of the fused grating. Five contrast ratios were tested for baseline. Three contrast ratios were used for post-patching measurement. D) Dichoptic masking. The subjects were asked to detect in which of two intervals the detection grating appeared. Two types of dichoptic mask were used. The parallel mask had the same orientation as the target. The cross-oriented mask had an orthogonal orientation.

**Figure 6:**
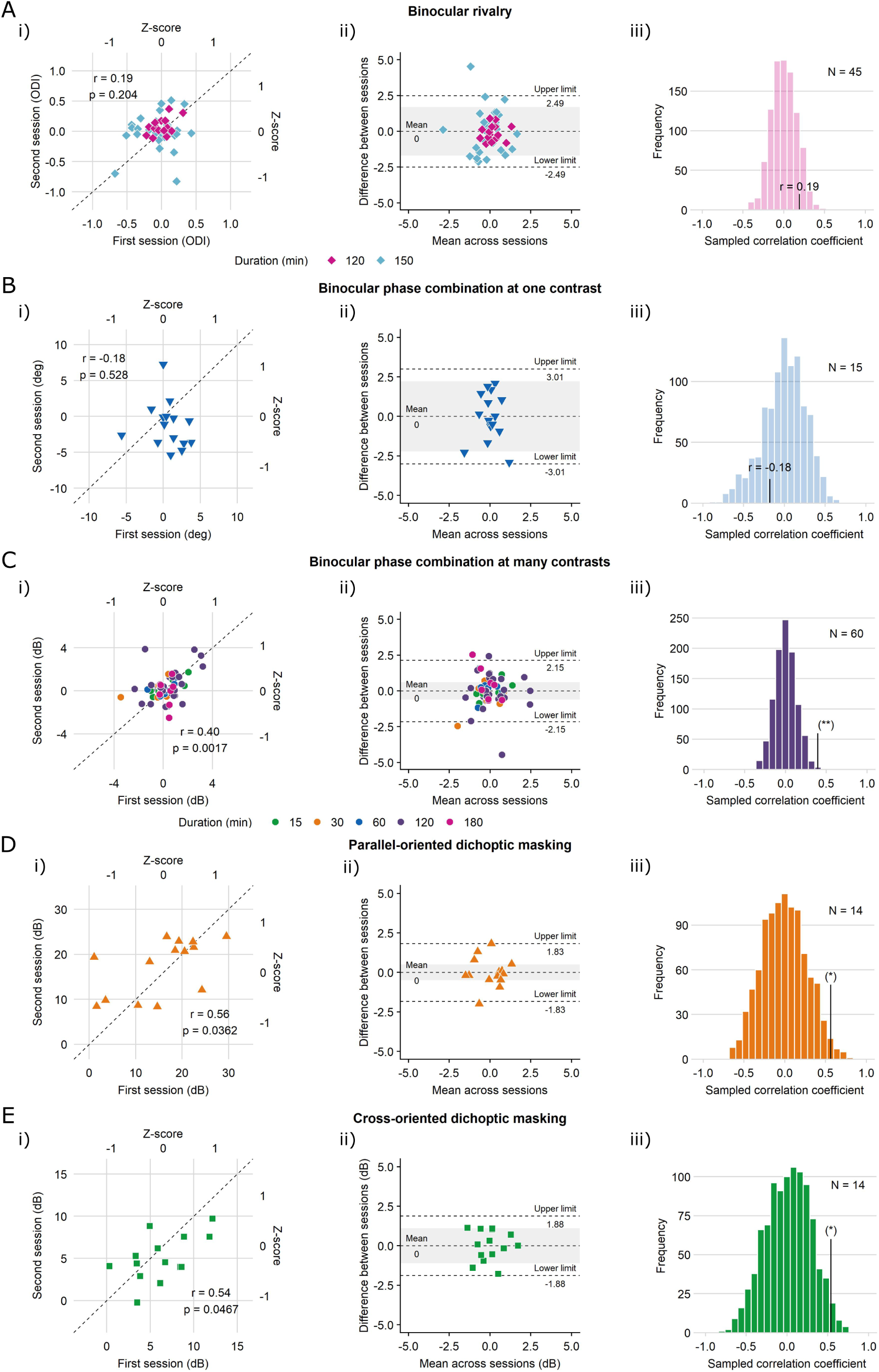
Evaluation of baseline measurement with the five psychophysical task variations.

#### 3.1.1 Binocular Rivalry

In binocular rivalry, ocular dominance index (ODI) indicates which of the percepts (patched or non-patched eye) shown separately to the both eyes dominate throughout the test block.

First, we investigated whether binocular rivalry is a reliable tool to study ocular dominance plasticity. To begin with, Pearson’s correlation was calculated to assess whether the baseline performance of a subject in one day is correlated to that of the same subject from another day. The correlation was not significant (n = 45, r = 0.19, p = 0.204, see Figure 6A(i)). Next, the raw data of ocular dominance index were converted into z-scores. All points except one seem to reside within the range of z-scores ± 1. This indicates that most points are within 1 standard deviation from the mean of the dataset for each session.

To see if there was a good agreement between the two experimental sessions, we created a Bland-Altman plot. Figure 6A(ii) indicates that the limits of agreement are ± 2.49 (z-scores). The limits of agreement (dashed lines) represent the test-retest variability that originate from multiple factors, such as day-to-day variability between the two experimental sessions and the inherent variability from the psychophysical measurement itself. Therefore, we computed the binocular rivalry measurement variability. This range is shown as a grey shaded area in Figure 6B, it is ± 1.69 (z-scores). The bulk of the area within the limits of agreement is taken up by the shaded region. This suggests that most of the test-retest variability originates from the binocular rivalry measurement itself, rather than variability in physiological factors.

Lastly, we evaluated whether the performance of a subject from the first experimental session was more correlated to that same subject’s performance from the second experimental session rather than that from another, randomly selected subject altogether. The sampled correlation coefficients are plotted in histogram (see Figure 6A(iii). As we expected from Figure 6A(i), the correlation between the performance scores in both experimental sessions is weak. This means that the measurement variability is so large that there is little to be gain from using a within-subject protocol to make comparisons.

#### 3.1.2 Binocular Combination at One Contrast

Pearson’s correlation was conducted for results for the same subjects on different days. They were not correlated (n = 15, r = −0.18, p = 0.528). All points except two points reside within the range of z-scores ± 1. The Bland-Altman plot is shown in Figure 6B(ii). The limits of agreement are ± 3.01 (z-scores). We calculated the measurement variability expected from the binocular combination measurement. This is shown as a grey shaded area in Figure 6B(ii), spanning ± 2.22 (z-scores). Since the shaded area makes up most of the area within the limits of agreement (dashed lines), most of the test-retest variability originates from the task measurement variability rather than from other factors.

The sampled correlation coefficients are plotted in histogram (Figure 6B(iii)). As expected from Figure 6B(i), the correlation between the performance scores in both sessions is weak. The correlation coefficient obtained from Figure 6B(i) falls within that of the histogram, suggesting that with this degree of measurement error, within-subject designs offer little if any advantage.

#### 3.1.3 Binocular Combination at Many Contrasts

Recently, a more extensive version of the binocular phase combination task has been used to study ocular dominance plasticity [4, 21, 28]. This version makes measurements at multiple contrast ratios and calculates the shift in ocular dominance using a model. For this data analysis, we treated the performance of one subject for a certain patching duration as a distinct data point from that of the same subject for another patching duration. As a result, there were 60 unique data points.

To begin with, Pearson’s correlation was calculated (see Figure 6C(i)). The correlation was significant (n = 60, r = 0.40, p < 0.01). All points point reside within the range of z-score 1. A Bland-Altman plot is illustrated in Figure 6C(ii), which indicates that the limits of agreement are ± 2.15 (z-scores). We computed the measurement variability from this binocular combination task. This range (shown as a grey shaded area in Figure 6C(ii)) is ± 0.61 (z-scores). The shaded area only represents a small fraction of the area within the limits, suggesting that most of the test-retest variability originates from external factors such as day-to-day variability in the physiological mechanisms. Lastly, the sampled correlation coefficients are plotted in a histogram (Figure 6C(iii)). As observed in Figure 6C(i), the correlation between the performance scores in both experimental sessions is robust. This is confirmed in Figure 6C(iii) where the correlation coefficient obtained from Figure 6C(i) resides outside that of the histogram. This suggests there is much to be gained from using within-subject testing protocols.

#### 3.1.4 Parallel-Oriented Dichoptic Masking

Baldwin and Hess (2018) used dichoptic masking to study the patching effect [20]. The magnitude of the patching effect was found to depend on the orientation of the masks. For this reason, we analysed data from the two tasks separately: parallel-oriented dichoptic masking and cross-oriented dichoptic masking.

For parallel-oriented dichoptic masking, Pearson’s correlation test revealed a significant correlation (n = 14, r = 0.56, p < 0.05; Figure 6D(i)). All points reside within the range of z-scores 1. A Bland-Altman plot is presented in Figure 6D(ii). The limits of agreement are ± 1.83 (z-scores). The measurement variability expected from the task alone is shown as a grey shaded area in Figure 6D(ii); its range is ± 0.50 (z-scores). The shaded area only represents a small fraction of the area within the limits of agreement. This suggests that most of the test-retest variability originates from external factors such as day-to-day variability. Lastly, the sampled correlation coefficients are plotted in histogram form (see Figure 6D(iii)). As we observed in Figure 6D(i), the correlation between the performance scores in both experimental sessions is robust. This is confirmed in Figure 6D(iii) where the correlation coefficient obtained from Figure 6D(i) seems to reside in the outer edge of the histogram. Therefore, there is an advantage to be had from within-subject testing protocols.

#### 3.1.5 Cross-Oriented Dichoptic Masking

For cross-oriented dichoptic masking, the Pearson’s correlation test was significant (n = 14, r = 0.54, p < 0.05; Figure 6E(i)). Next, the raw data of grating threshold (dB) were converted into z-scores. All points except one reside within the range of z-scores ± 1. The Bland-Altman plot is shown in Figure 6E(ii). This shows the limits of agreement are ± 1.88 (z-scores). The expected measurement variability arising from the dichoptic masking task itself is shown as a grey shaded area in Figure 6E(ii). The range of the shaded area is ± 1.10 (z-scores). It seems the larger portion of the areas within the limits of agreement are attributable to the measurement variability from the dichoptic masking task itself rather that from external factors such as day-to-day variability. However, it is notable that the additional area within the limits of agreement that is attributable to external factors is of a similar size. Lastly, the sampled correlation coefficients are plotted in histogram (see Figure 6E(iii)). As we observed in Figure 6E(i), the correlation between the performance scores in both experimental sessions is strong. This is confirmed in Figure 6E(iii) where the correlation coefficient obtained from Figure 6E(i) resides at the outer edge of the histogram, suggesting that within-subject testing protocols are advantageous.

Figure 6 is divided into five rows (task) and three columns (as described in the Standardized Data Analysis section). Row (A) Binocular rivalry. Pink points represent data from the new experiments that implemented a patching duration of 120 minutes (n = 15), blue points from the study of Finn et al. (2019) where subjects were patched for 150 minutes (n = 30). Row (B) Binocular phase combination at one contrast. 15 subjects were patched for 120 minutes. Row (C) Binocular phase combination at many contrasts. Different durations of patching are represented in different colors. Row (D) Parallel-oriented dichoptic masking. 14 subjects were patched for 150 minutes. Row (E) Cross-oriented dichoptic masking. 14 subjects were patched for 150 minutes. Column (i) Baseline reliability. The x-axis represents ocular dominance index from the first experiment session, and the y-axis from the second session. The secondary x- and y-axes represent z-scores from the raw data of ocular dominance index. The dashed line represents the line of equality (1st session = 2nd session). Each diamond represents a data point of one subject. Column (ii) Baseline measurement variability illustrated in a Bland-Altman plot. Difference in z-scores between the first and second session is plotted as a function of the mean of z-scores across two sessions. The outer horizontal dashed lines indicate 95% limits of agreement. The dashed line in the middle indicates the mean difference of z-scores across the subjects. The gray shaded region within the limits of agreement represent baseline consistency (i.e., the testing variability stemming from only the binocular rivalry task). The unshaded regions within the limits of agreement represent test-retest variability from external factors beside the task itself. Column (iii) Baseline reliability illustrated in a histogram. The sampled reliability coefficients are plotted as a histogram, where the y-axis represents the frequency and the x-axis the sampled correlation coefficient ranging from −1 to 1. The single line value represents the within subject correlation and this is compared to the distribution of across subjects correlations.

### 3.2 Magnitude of Changes in Sensory Eye Balance after Short-Term Patching

#### 3.2.1 Binocular Rivalry

The patching effect is represented by the difference in ocular dominance index between baseline and post-patching measurements. The more positive the Δ ODI, the stronger the patching effect. A Pearson’s correlation test revealed a non-significant correlation (n = 15, r = 0.15, p = 0.597). All points reside within the range of z-scores ± 1.

The Bland-Altman plot in Figure 7A(ii) indicates that the limits of agreement are ± 2.56 (z-scores). The measurement variability from the binocular rivalry task itself is shown as a grey shaded area in Figure 7A(ii). Its range is ± 1.49 (z-scores). Similar to in the baseline measurements, the shaded area makes up the bulk of the area within the limits of agreement. This suggests that most of the test-retest variability of the patching effect originates from the measurement error of the binocular rivalry task itself.

**Figure 7:**
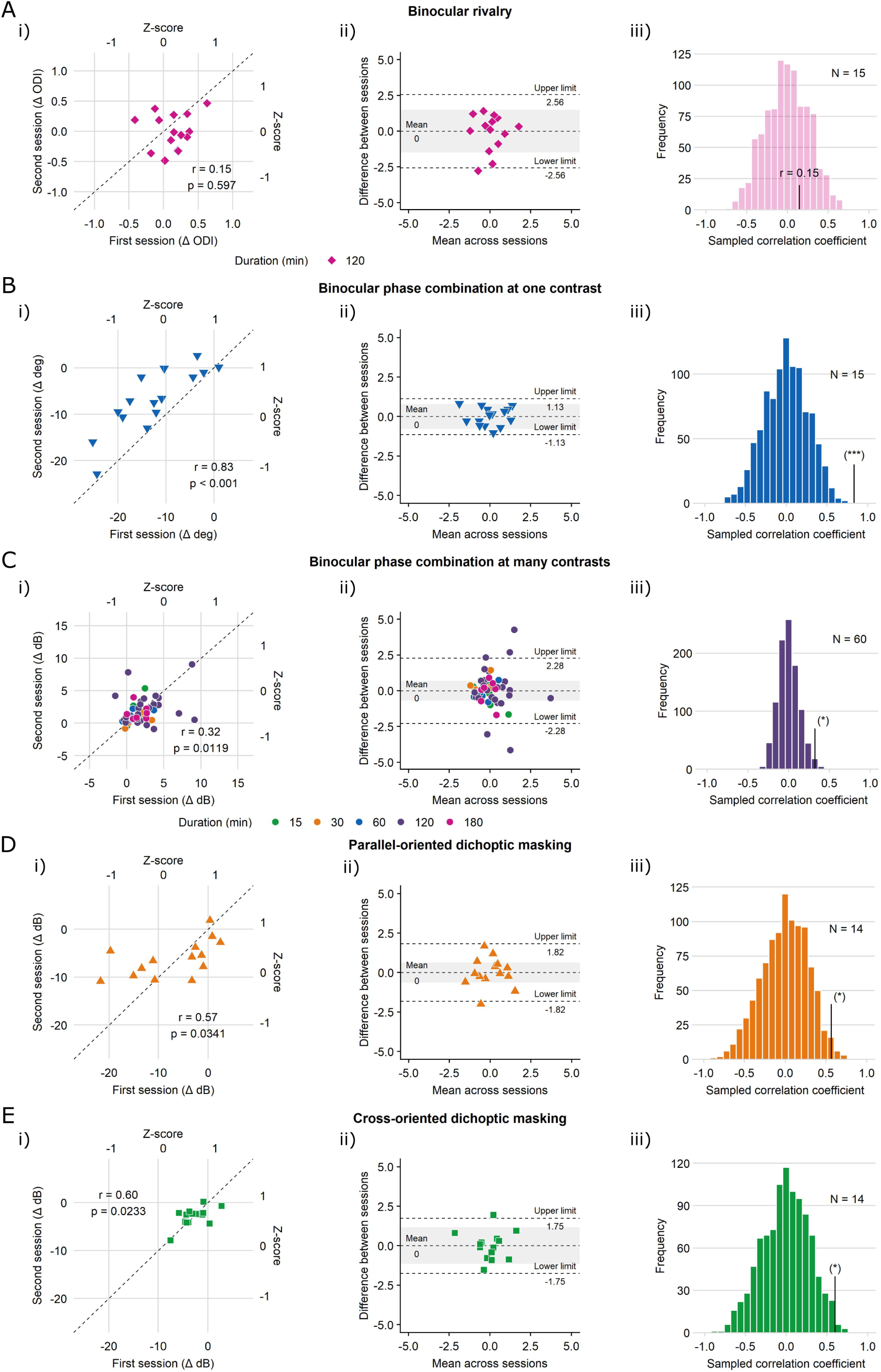
Repeatability of the patching effect as measured in the five psychophysical task variations.

Histogram of the sampled correlation coefficients are plotted Figure 7A(iii). The correlation between the patching effect scores in both experimental sessions is weak (see Figure 7A(i)). This is confirmed in Figure 7A(iii) where the correlation coefficient obtained from Figure 7A(i) resides in the middle of the histogram, suggesting that it is not beneficial to use subjects as their own control.

#### 3.2.2 Binocular Combination at One Contrast

The change in sensory eye dominance from patching is represented by the difference in perceived phase (deg) between baseline and post-patching measurements. The more negative the difference in perceived phase, the stronger the patching effect). A Pearson’s correlation test found a significant correlation (n = 15, r = 0.83, p < 0.001). All points except two reside within the range of z-scores ± 1. The Bland-Altman plot in Figure 7B(ii) indicates that the limits of agreement are ± 1.13 (z-scores). The expected measurement variability from the binocular combination task itself is illustrated as a grey shaded area in Figure 7B(ii); its range is ± 0.80 (z-scores).

Histogram of the sampled correlation coefficients are plotted Figure 7B(iii). The correlation between the patching effect scores in both experimental sessions is very robust (see Figure 7B(iii)). This is confirmed in Figure 7B(iii) where the correlation coefficient obtained from Figure 7B(i) is located outside the histogram, suggesting that within-subjects designs are advantageous.

#### 3.2.3 Binocular Combination at Many Contrasts

The change in sensory eye dominance from short-term patching is represented by the difference in contrast ratio (Δ dB) between baseline and post-patching measurements. The more positive the difference in contrast ratio (Δ dB), the stronger the patching effect. Results from a Pearson’s correlation test were significant (n = 60, r = 0.32, p = 0.012; Figure 7C(i)). All points are within the range of z-scores ± 1.

The Bland-Altman plot in Figure 7C(ii) indicates that the limits of agreement are ± 2.26 (z-scores). All points except five are within the limits of agreement. The expected measurement variability from the binocular combination task itself is presented in Figure 7C(ii) as a grey shaded area; its range is ± 0.70 (z-scores). Most of the area within the limits of agreement is not shaded in grey. That means most of the test-retest variability from the patching effect originates from factors other than the measurement variability associated with binocular combination task itself.

Histogram of the sampled correlation coefficients are plotted Figure 7C(iii). The correlation between the patching effect scores in both experimental sessions is robust (see Figure 7C(i)). This is confirmed in Figure 7C(iii) where the correlation coefficient obtained from Figure 7C(i) resides in the outer edge of the histogram, suggesting that within-subjects designs are more sensitive than between-subject designs.

#### 3.2.4 Parallel-Oriented Dichoptic Masking

The change in sensory eye dominance from patching is represented by the difference in contrast ratio (dB) between baseline and post-patching measurements. The more negative the difference in the contrast threshold for the test grating (Δ dB), the stronger the patching effect. This applies to both parallel- and cross-oriented dichoptic masking. A Pearson’s correlation test revealed a significant correlation (n = 14, r = 0.57, p < 0.05; Figure 7D(i)). All points except one reside within the range of z-scores ± 1.

The Bland-Altman plot in Figure 7D(ii) indicates that the limits of agreement are ± 1.82 (z-scores). All points except one are within the limits of agreement. The expected measurement variability from the task is shown in Figure 7D(ii) as a grey shaded area, which has a range of ± 0.64 (z-scores). Most of the area within the limits of agreement is not shaded in grey. This indicates that most of the test-retest variability of the patching effect originates from factors other than the task measurement error. Histogram of the sampled correlation coefficients are plotted Figure 7D(iii). The correlation between the patching effect scores in both experimental sessions is robust (see Figure 7D(i)). This is confirmed in Figure 7D(iii) where the correlation coefficient obtained from Figure 7D(i) resides in the outer edge of the histogram, suggesting that within-subject designs are superior to between-subject designs.

#### 3.2.5 Cross-Oriented Dichoptic Masking

A Pearson’s correlation test indicated a significant correlation (n = 14, r = 0.60, p < 0.05; Figure 7E(i)). All points except one reside within the range of z-scores ± 1.

The Bland-Altman plot in Figure 7E(ii) indicates that the limits of agreement are ± 1.75 (z-scores). All points except one are within the limits of agreement. The expected measurement variability from the task itself is shown as a grey shaded area in Figure 7E(ii); its range is 1.16 (z-scores). Most of the area within the limits of agreement is shaded in grey. This suggests that most of the test-retest variability of the patching effect originates from the task measurement itself. Histogram of the sampled correlation coefficients are plotted Figure 7E(iii). The correlation between the patching effect scores in both experimental sessions is robust (see Figure 7E(i)). This is confirmed in Figure 7E(iii) where the correlation coefficient obtained from Figure 7E(i) resides in the outer edge of the histogram, suggesting that there is an advantage of using a within-subject design for this task.

Figure 7 is divided into five rows (task) and three columns (data analyses). Row (A) Binocular rivalry. 15 subjects were patched for 120 minutes. Row (B) Binocular phase combination at one contrast. 15 subjects were patched for 120 minutes. Row (C) Binocular phase combination at many contrasts. Different durations of patching are represented in different colors. Row (D) Parallel-oriented dichoptic masking. 14 subjects were patched for 150 minutes. Row (E) Cross-oriented dichoptic masking. 14 subjects were patched for 150 minutes. The columns present data in the same manner as in Figure 6.

### 3.3 Summary of Results

We have evaluated and compared four properties for five variants of psychophysical task, each having been used in the past to assess the patching effect. These properties are *baseline reliability, patching effect reliability, baseline measurement variability and patching effect measurement variability* (defined in the Introduction). The correlations (i.e. reliabilities) for baseline measurements and for the magnitude of the patching effect are summarised as p-values from Pearson’s correlation tests between the raw data from the first and second experimental sessions. The baseline and the patching effect measurement variabilities are summarised as the measurement error from the psychophysical task itself rather than extraneous errors such as day-to-day variability. As we previously mentioned, baseline data provide the reliability and repeatability of the task because confounds such as visual deprivation have been removed. The only effects are day to day physiological or psychological variation, and the variability in the measurement from the task itself. On the other hand, the magnitude of the patching effect was quantified by the magnitude of change in sensory eye dominance after patching relative to baseline. Therefore, it includes the baseline effects and also any variability in the strength of the patching effect across days.

In order to rank the psychophysical tasks from best to worst, we normalised the statistical values that represent each of the four pivotal properties across all tasks using the equation:

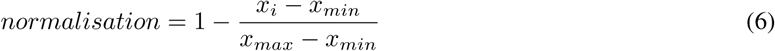

where *x*_*i*_ indicates a value that ought to be normalised within the dataset (e.g., p = 0.204 in baseline correlation/reliability of binocular rivalry), *x*_*min*_the minimum value in the dataset, and *x*_*max*_ the maximum value in the dataset. If the normalised value were 1, it would indicate that it was the best; if the normalised value were 0, it would indicate that it was the worst. In the case of the reliabilities of baseline and the patching effect, the lowest p-value from Pearson’s correlation tests across the tasks was converted to 1, and the highest p-value to 0. However, for the measurement variabilities of baseline and the patching effect, the smallest standard error within the limits of the agreement (grey areas from columns (ii) in Figures 6 and 7) was converted to 1, and the widest range to 0. Figure 8 shows the summary ranking of these four properties.

**Figure 8:**
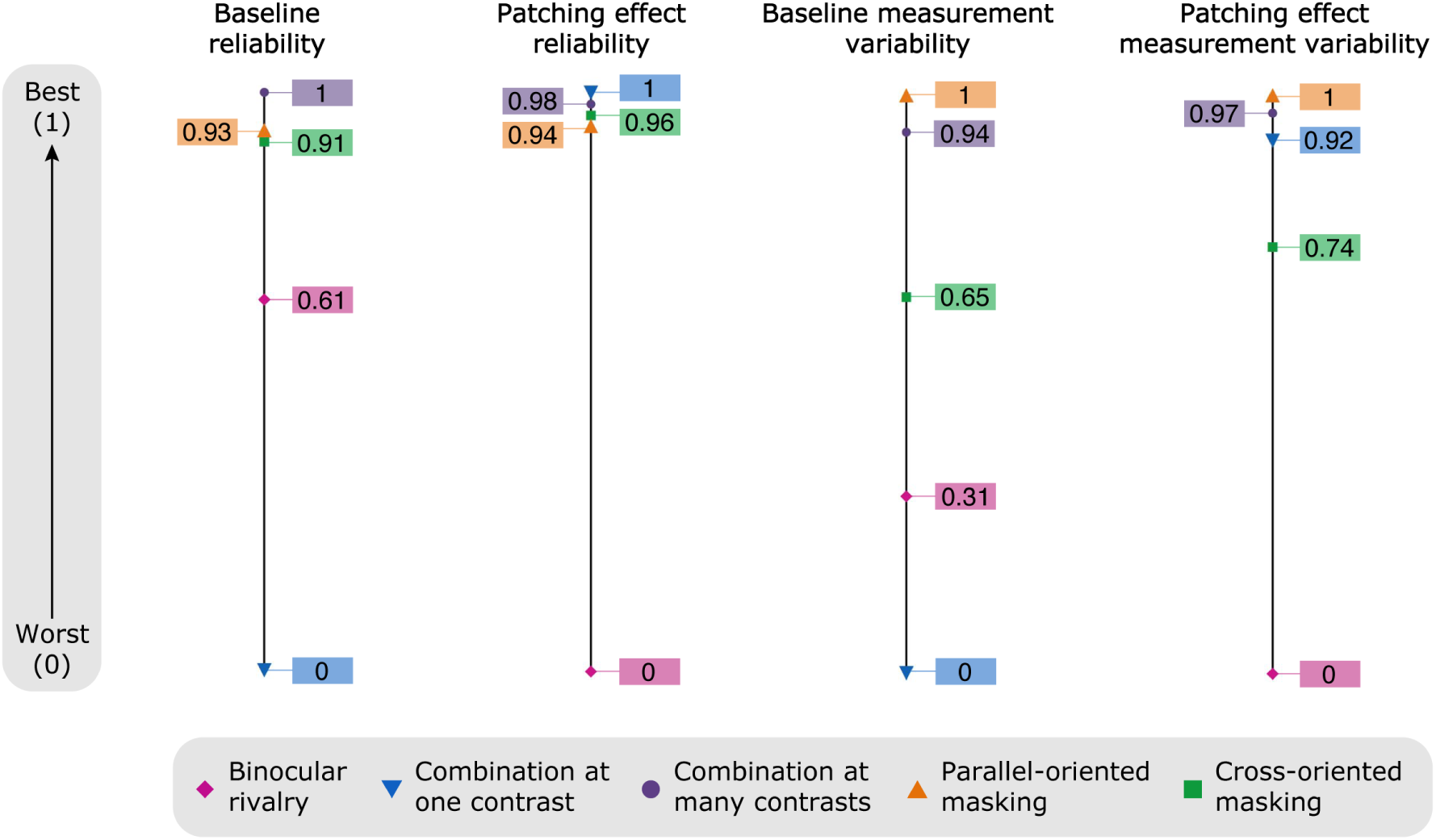
Summary of results. Each value was normalised in a scale where 1 represents best and 0 worst of all tasks.

## 4 Discussion

### 4.1 Are Different Psychophysical Tasks Associated with Distinct Neural Sites and Mechanisms?

Studies using binocular rivalry and binocular combination at one contrast have revealed differences in the magnitude of the patching effect [6, 14, 15]. This leads to the notion that the patching effect is a phenomenon that occurs at multiple neural sites. If this interpretation is true, different psychophysical tasks might be associated with different aspects/sites. However, we show that this difference in results could also be attributed to a wide measurement variability of the patching effect as demonstrated in binocular rivalry (see Figure 8). Whether the discrepancies between the two psychophysical tasks are purely due to the principle that the patching effect is multifaceted, or the wide measurement variability of the tasks remains to be resolved.

On the other hand, Baldwin and Hess (2018) used dichoptic masking with two orientations of the mask grating to emulate binocular combination and rivalry within the same task [20]. This choice enabled them to remove measurement variability from using multiple psychophysical tasks. They reported that the orientation of the mask determined the magnitude of the patching effect. This finding reinforces the notion that the patching effect is multifaceted, and that one psychophysical task might capture only one aspect of the neural plasticity change.

### 4.2 How Reliable is Baseline Measurement for Each Task?

As Figure 8 shows, binocular rivalry and binocular combination at one contrast have poor reliability and measurement variability in baseline measurement, whereas binocular combination at multiple contrasts and dichoptic masking at both orientations seem to measure baseline reliability. What could be contributing factors for the poor reliability of binocular rivalry and binocular phase combination at one contrast?

First, the binocular rivalry task has been used to study a wide range of visual phenomena [29]. It represents the competition, rather than the combination, between the eyes by presenting two rivalrous images separately to both eyes. The interocular competition during rivalry causes a rapid and irregular fluctuation of sensory eye dominance over visual space and time [30, 29, 31]. The random nature of binocular rivalry might have contributed to the large measurement variability of the baseline. Moreover, attention can affect the temporal dynamics of rivalry [32], suggesting that this task is significantly influenced by cognitive factors. The poor reliability of baseline measurement between the two separate days of testing might indicate that the level of attention throughout the task between the sessions was dissimilar. Therefore, it seems that the random dynamic nature of binocular rivalry and the influence of top-down attentional factors could have contributed to inducing a large measurement variability for baseline measurement.

Another explanation for the poor baseline measurement is that only one contrast ratio between the eyes was used. In binocular combination at many contrasts, the subjects were tested at various contrast ratios instead. Subsequently, the contrast ratio where the perceived phase is 0 was estimated by fitting a contrast gain model [23] to the data across all contrast levels. Therefore, the version of the task in which more data is collected across multiple contrast values, not surprisingly, has much tighter measurement variability.

### 4.3 Is the Patching Effect Stable Across Days?

For all five psychophysical task variations, most subjects experienced a significant shift in eye dominance in favour of the patched eye, a phenomenon that we would expect from short-term patching in adults. Therefore, the patching effect is repeatable across tasks. Indeed, most studies using various tests have demonstrated the replicability of the patching effect. However, the magnitude of the plasticity change does not seem to be uniform across tasks. In addition, it seems to be notably different across separate days using the same task. For instance, if the patching effect is measured with the binocular rivalry task, it is not well correlated across separate days (poor reliability; see Figure 8). As is the case in baseline performance, the irregular dynamics and attentional factors associated with binocular rivalry over space and time could explain the poor repeatability of the patching effect [30, 29, 31, 32]. On the other hand, the other four psychophysical tasks seem to show a similar magnitude of the patching effect across days, as demonstrated in the tight measurement variability of the patching effect. Moreover, the reliability of the patching effect across days for all tasks is demonstrated by robust correlations. Therefore, the four tasks show a high repeatability of the patching effect. These findings refute the argument that the patching effect fluctuates across days within the same subject and advocate that the variability of the patching effect stems depends on the task.

In addition, it is noteworthy to compare the Bland Altman plots of rows C-E of Figures 6 and 7. The limits of agreement and measurement variability appear similar between the two figures. This is important, as it indicates that there is not a further additional source of variability introduced by patching. This evidence also rebuts the idea that the patching effect is itself inherently variable across days.

### 4.4 Which psychophysical tasks should be used in the clinical setting to measure the patching effect?

Recent clinical studies on amblyopes have incorporated training protocols that involve patching the dysfunctional eye [33, 34, 35], a design that is identical to the one used in short-term patching studies in controls. Therefore, the choice of test for measuring the patching effect might also guide the development of clinical treatment. To ensure that the findings from preliminary studies are replicable in a wider population, the choice of test in clinical studies is important. Our findings show that binocular rivalry and binocular combination at only one contrast are not ideal tasks. Binocular rivalry seems to be particularly variable for measuring the patching effect. This may limit its utility for clinical studies. Instead, we recommend psychophysical tasks that best capture stable baseline performance and a repeatable patching effect. According to our results, these tasks are binocular phase combination at multiple contrasts and parallel-oriented dichoptic masking.

## 5 Conclusion

There have been conflicting reports on the patching effect as a result of short-term deprivation in adults and children. The magnitude of the patching effect has been variable across different tests (binocular rivalry and combination) and within the identical test (binocular rivalry) across conditions. In the Introduction, we proposed three explanations for these discrepancies. First, the mechanism of this patching effect might be multifaceted and different tasks might reveal different aspects/sites. If this notion holds true, each psychophysical task might capture only one aspect of the entire plasticity change. Previous psychophysical studies have shown this to be the case [6, 20]. Second, the measurement error associated with the tasks might be poor. In light of our findings, this claim seems to be a reasonable explanation for some tasks. In addition to showing that binocular rivalry and combination at one contrast show poor repeatability of baseline performance, our study shows that binocular rivalry exhibits a poor repeatability for the magnitude of the patching effect. Third, the patching effect might be itself an unstable phenomenon. Our findings show that this is not the case, as we do not find evidence of additional variability in the measurements made when subjects had been patched.

## 6 Acknowledgment

This work was supported by the National Natural Science Foundation of China (31970975), the Qianjiang Talent Project (QJD1702021), the Wenzhou Medical University grant QTJ16005 and the Project of State Key Laboratory of Ophthalmology, Optometry and Visual Science, Wenzhou Medical University (K171206) to JZ, the Zhejiang Basic Public Welfare Research Project (LGJ20H120001) to ZH, the Canadian Institutes of Health Research Grants CCI-125686, NSERC grant 228103, and an ERA-NET Neuron grant (JTC2015) to RH, and Canadian institutes of Health Research graduate award to SM. The sponsor or funding organization had no role in the design or conduct of this research. This pre-print manuscript was written in LATEX using Overleaf. It is formatted with a custom style available at: github.com/alexsbaldwin/biorxiv-inspired-latex-style

